# Does genetic variation in controlled experiments predict phenology of wild plants?

**DOI:** 10.1101/2024.09.02.610887

**Authors:** Victoria L. DeLeo, David L. Des Marais, Claire M. Lorts, Thomas E. Juenger, Jesse R. Lasky

## Abstract

Phenology and the timing of development are often under selection. However, the relative contributions of genotype, environment, and prior developmental transitions to variance in the phenology of wild plants is largely unknown. Individual components of phenology (e.g., germination) might be loosely related with the timing of maturation due to variation in prior developmental transitions. Given widespread evidence that genetic variation in life history is adaptive, we investigated to what degree experimentally measured genetic variation in Arabidopsis phenology predicts phenology of plants in the wild. As a proxy of phenology, we obtained collection dates from nature of 227 naturally inbred *Arabidopsis thaliana* accessions from across Eurasia. We compared this phenology in nature with experimental data on the descendant inbred lines that we synthesized from two new and 155 published controlled experiments. We tested whether the genetic variation in flowering and germination timing from experiments predicted the phenology of the same lines in nature. We found that genetic variation in phenology from controlled experiments significantly predicts day of collection from wild individuals, as a proxy for date of flowering, across Eurasia. However, local variation in collection dates within a region was not explained by genetic variance in phenology in experiments, suggesting high plasticity across small-scale environmental gradients or complex interactions between the timing of different developmental transitions. While experiments have shown phenology is under selection, understanding the subtle environmental and stochastic effects on phenology may help to clarify the heritability and evolution of phenological traits in nature.

## Introduction

The predictability of trait variation from genetic variation, *i.e.* trait heritability, is a key quantity determining the rate of trait evolution and the ability of populations to adapt to changes in selection (Lush 1937; Gomulkiewicz and Houle 2009). Plant biologists have long relied on common garden experiments to estimate breeding values and heritability for a range of traits (Mazer and LeBuhn 1999). However, natural populations often occur in highly heterogeneous environments to which organismal traits may respond plastically. As a result, heritability in natural populations may be much lower than those in common gardens. Furthermore, even field common gardens might fail to accurately recreate natural selective pressures if the direction and magnitude of selection varies in time (Karrenberg & Widmer 2008). As an alternative to common gardens, heritability can be estimated using pedigrees or genomic data from wild individuals (Gienapp et al. 2017). However, these models rely on assumptions of certain genetic architectures (Mitchell-Olds and Rutledge 1986) and the resulting heritability estimates are not often compared with common garden estimates of trait variation (but see (Weigensberg and Roff 1996)).

Phenology, or the timing of the developmental transitions between an organism’s life stages, directly and indirectly influences plant fitness and selection by determining the traits that are expressed at any point throughout the year (Donohue 2005). In seasonally variable environments, traits such as flowering time and dormancy can ameliorate harsh abiotic conditions by timing dormant periods to coincide with unfavorable seasons (e.g. drought escape Ludlow 1989; Lawrence-Paul and Lasky 2024). Flowering, vegetative phase change, and germination timing often show geographic clines, suggesting their importance in adaptation to local environments (Baughman et al. 2019; Martínez-Berdeja et al. 2020; Lawrence-Paul et al. 2023, 2025). Common garden studies often find that genetic variation in individual components of phenology are under selection and have further demonstrated variable strength and direction of selection across field sites (Hall and Willis 2006; Fournier-Level et al. 2013; Ågren et al. 2017; Gamba et al. 2024b). However, the duration of individual developmental transitions (e.g. flowering time) might be loosely related to phenology in nature due to variation in prior developmental transitions (e.g. germination) (Burghardt et al. 2015). While phenology is determined by both genetic factors and plastic responses to environment (Amasino 2004; Andrés and Coupland 2012; Auge et al. 2018), the relative importance of these two sources of phenotypic variance in the timing of individual developmental transitions determining phenology in nature is unclear, even in model systems.

Much of our understanding of the genetic basis of plant phenology comes from studies in the model *Arabidopsis thaliana* (hereafter Arabidopsis). Common garden trials in both lab and field settings have uncovered genetic loci contributing to much of the observed variation in dormancy and flowering time (Juenger et al. 2005; Atwell et al. 2010; Brachi et al. 2010; Fournier-Level et al. 2013; Ågren et al. 2017). Flowering time and seed dormancy are determined by complex, overlapping gene networks, leading to pleiotropy involving these traits (Simpson and Dean 2002; Wilczek et al. 2010; Auge et al. 2019). These traits are also plastic and show genotype by environment interactions (Juenger et al. 2005; Zhou et al. 2005; Wilczek et al. 2009; Lorts and Lasky 2020): germination responds to temperature, photoperiod, moisture, and nutrient availability (Huang et al. 2010, 2018; Penfield and Springthorpe 2012; Footitt et al. 2013; Kenney et al. 2014), while flowering responds to multiple cues such as temperature and daylength (Thomas and Vince-Prue 1997; Lempe et al. 2005; Balasubramanian et al. 2006).

The broad phenological variation described in controlled environment experiments is observed in wild populations as well (Ratcliffe 1976; Simpson and Dean 2002; DeLeo et al. 2020). Arabidopsis can exhibit the life history of a winter annual, germinating in the fall, overwintering as a rosette, and flowering in the spring (Figure 1). However, Arabidopsis can also germinate and flower in a single growing season. These shorter-lived plants tend to be facultative — flowering in spring, summer, or fall — and in some regions this rapid life cycle enables multiple generations within a year. This variation in life histories occurs in many annual plants (Baskin and Baskin 1988) and is made possible, in part, by a range of flowering times and germination traits that can vary independently from each other (Marcer et al. 2018; Martínez-Berdeja et al. 2020). In Arabidopsis, geographic clines in genetic values for phenology, clines in allele frequencies of phenology QTL, and evidence of selection on phenology QTL support the conclusion that phenology is adapted to local environmental conditions in the wild (Caicedo et al. 2004; Stinchcombe et al. 2004; Samis et al. 2012; Fournier-Level et al. 2013; Ågren et al. 2017; Debieu et al. 2018; Exposito-Alonso et al. 2018; Lasky et al. 2018, 2024; Gamba et al. 2024a) (Figure 1). Yet, even within geographic proximity one can find genotypes with substantial genetic differences in flowering time (Jones 1971; Alonso-Blanco et al. 2016; Méndez-Vigo et al. 2022).

**Figure 1.**
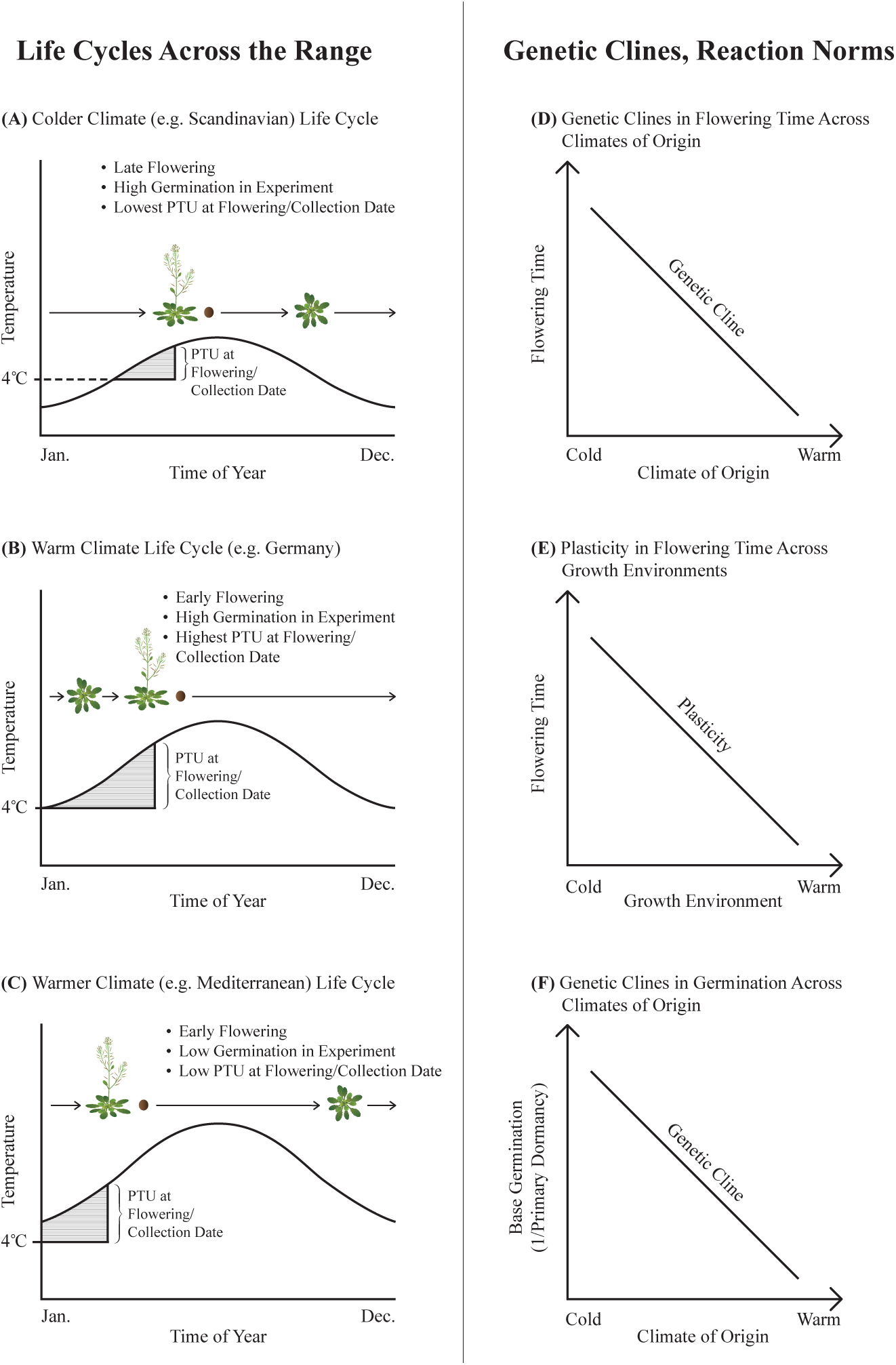
Genotype and environment likely influence phenology of individual Arabidopsis plants across its geographic range in the wild. Genetics and environmental cues may determine how long a plant remains in vegetative growth (green) or how long a plant remains dormant as a seed (brown circles), with three representative life cycles shown from Scandinavia to the Mediterranean (A–C), with shaded areas showing accumulation of photothermal units (PTUs) in the growing season (e.g. temperature >4°C). Existing knowledge of clines in flowering time and germination combined with plastic acceleration of flowering in warmer temperatures (D–F).

There are several mechanisms that may explain why phenology observed in controlled experiments shows poor correlation with phenology observed in nature (Wilczek et al. 2009). While genotype likely influences the phenology of an individual plant in the wild, the dependence of sequential developmental transitions on each other, as well as plasticity in their expression, may limit the translation of genetic values of single stage to natural phenology. First, there are interactions between germination and flowering time, both because of a shared genetic basis and because the timing of early life transitions influences environmental exposures in later life stages (Chiang et al. 2009; Huang et al. 2010; Springthorpe and Penfield 2015; Huo et al. 2016). Secondly, there is stochasticity in individual germination and flowering time (Jimenez-Gomez et al. 2011; Abley et al. 2020). Thirdly, spatial environmental variation is extensive in nature and genotype-environment interactions have major impacts on phenology (Sasaki et al. 2015). Finally, maternal effects on phenological transitions may be strong, as reported extensively for seed dormancy (Boyd et al. 2007; Chiang et al. 2013; Burghardt et al. 2016; Huang et al. 2018). As a result, flowering time loci identified in common garden lab or field experiments might have loose relationships with variation among natural individuals (Chiang et al. 2013); genotypes differing in phenology in one set of conditions may express the same phenology in another set of conditions (Wilczek et al. 2009; Burghardt et al. 2016).

Here, we compare phenological observations of wild individual plants from natural history collections (herbaria and seed stock centers) with plants in controlled environment experiments. Collection dates of natural history accessions have been demonstrated as accurate estimates of phenology (Miller-Rushing et al. 2006; MacGillivray et al. 2010; Davis et al. 2015; Ramirez-Parada et al. 2022). Across Eurasian Arabidopsis, we showed previously that herbarium collection dates were accelerated by approximately one week for each degree increase in spring temperatures and that there was a latitudinal cline in mean collection dates from early March to late June (DeLeo et al. 2020). Arabidopsis are naturally inbred (Jones 1971), thus we can compare collection dates of wild individuals with phenology of nearly genetically identical descendants in controlled experiments. This comparison provides a window into how genetic variation shapes phenology in natural populations against the forces of plasticity and GxE. Natural history collections can be used to characterize the climate conditions during development preceding collection (Bontrager et al. 2025b). We estimated phenology as collection date, and using the climate conditions preceding collection, we also estimated phenology as accumulated photothermal units (PTUs) at collection (Burghardt et al. 2015). PTUs allow us to account for weather effects on development and in essence allow estimation of how far into growing seasons plants were collected.

We developed three hypotheses for how the phenology of individual plants in nature may relate to genetic variation in phenological traits, illustrated in Figures 2-3:

- H1.We predict that Arabidopsis plants collected earlier in a natural growing season will have early flowering and low germination genetic values in controlled environments. We hypothesize that this pattern is due to co-gradient variation in flowering time along temperature gradients (i.e. when genetic clines and plastic responses to environment are positively correlated) (Levins 1968). Specifically, genetic clines in flowering time (here, early flowering in warmer climates) and germination rate, combined with plastic acceleration of flowering in warmer conditions (Figure 1), leads to range-wide positive relationships between genetic values for flowering time (Figure 2A), germination rate (Figure 2B), and collection date (as a proxy for flowering date in the wild).
- H2.For cumulative photothermal units (PTU) at collection, we predict that spring annuals with high germination rates and early flowering times will have been collected later in growing seasons (high PTU) than winter annuals with later flowering genetic values (Figure 2C, 2D).
- H3.Within local regions, environmentally induced phenological variation (i.e., plasticity associated with different timing of spring warming) is likely lower than that seen across the species range. Thus we predict that flowering time genetic values and germination rates would be positively related to local within-region variation in collection dates largely due to genetic effects (Figure 3).

**Figure 2:**
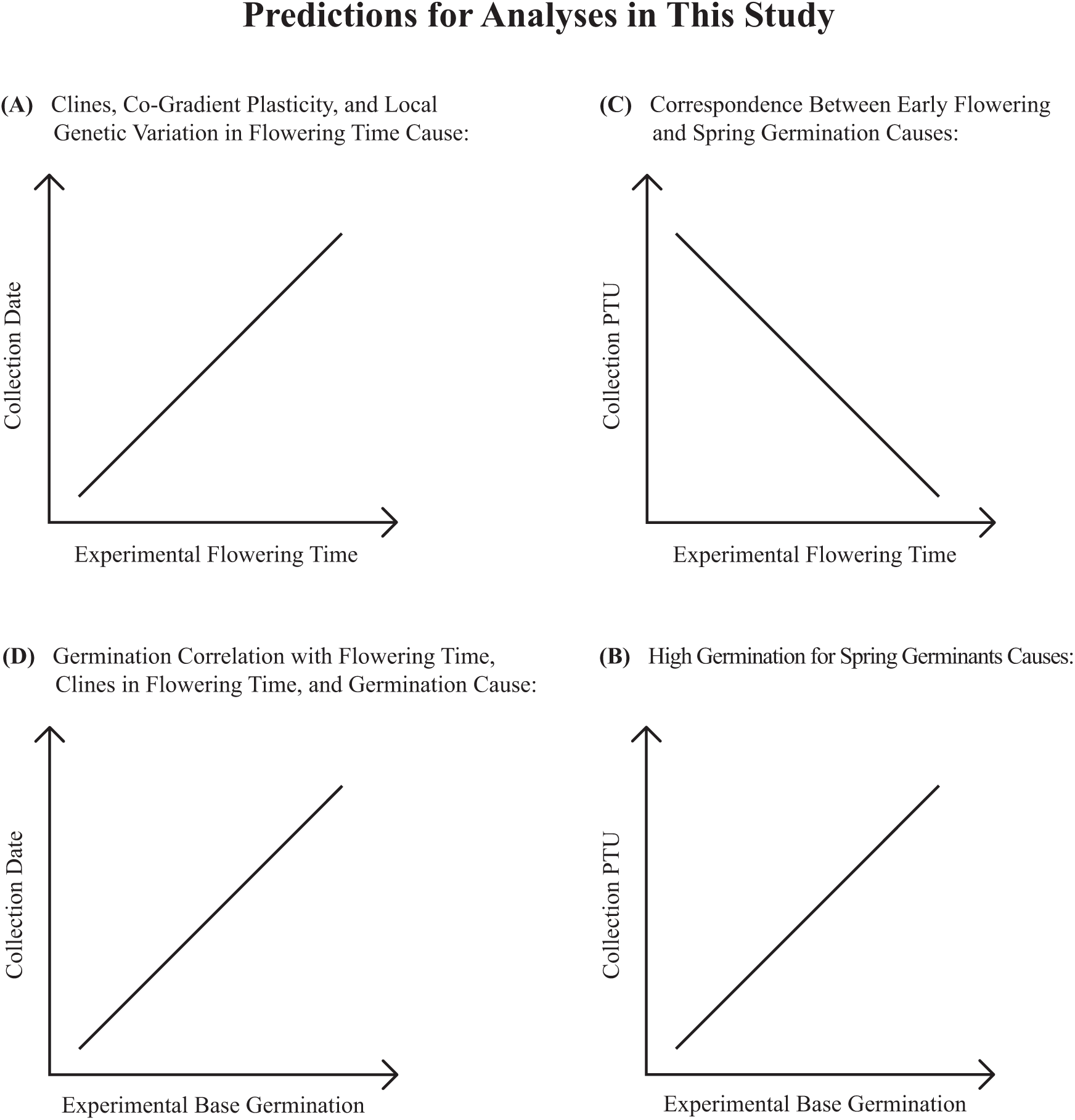
Existing knowledge of clines in flowering time and germination combined with plastic acceleration of flowering in warmer temperatures (see Figure 1) lead to our predictions (A–D here) of how flowering time and base germination rates in controlled experiments will correspond to phenology in natural wild plants.

**Figure 3.**
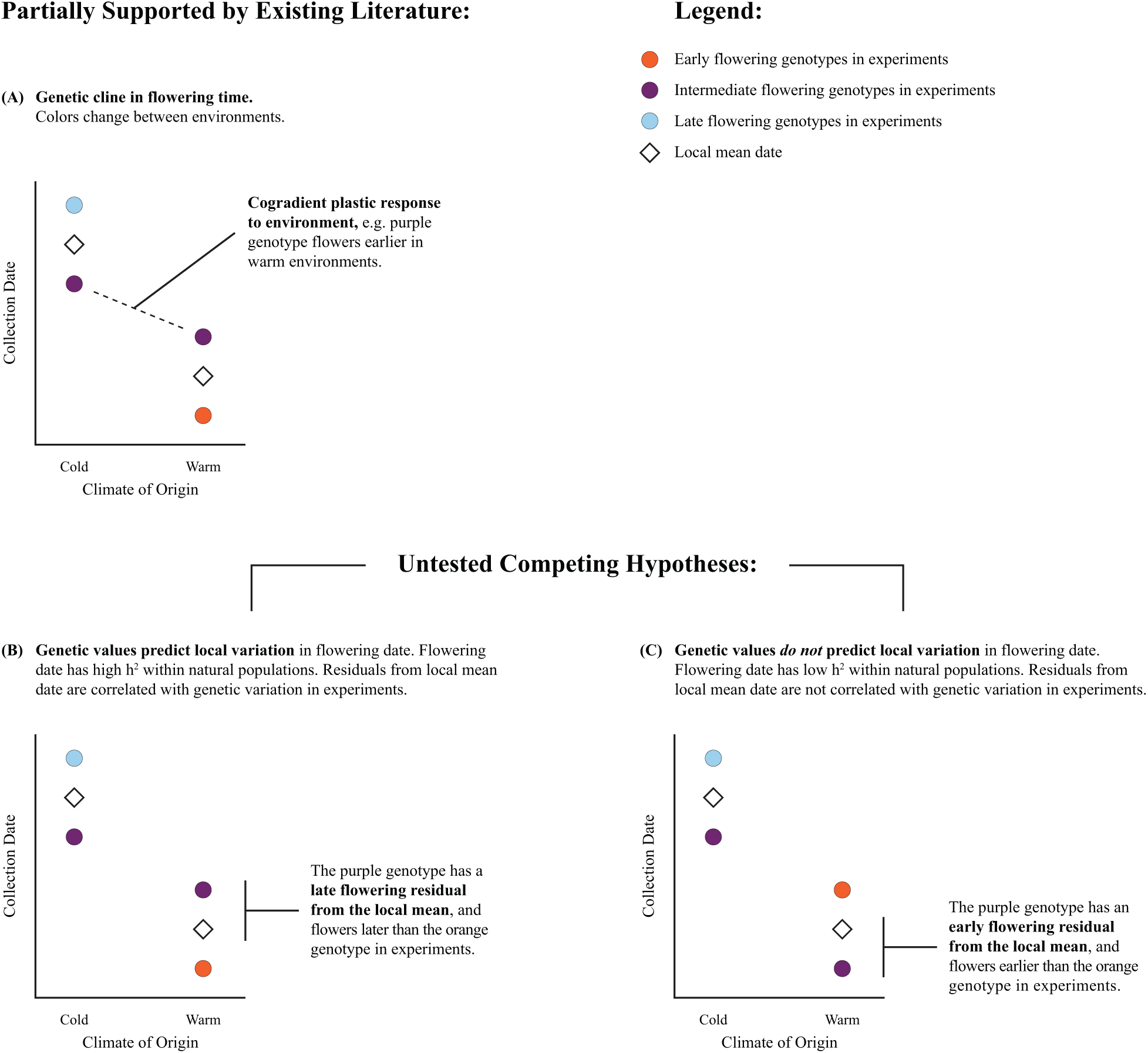
An illustration of hypotheses for local genetic variation in phenology within regions and the relationship with phenology of wild plants. In the presence of co-gradient variation (correlated plasticity and genetic clines, panel A) it is important to account for local mean collection dates (diamonds) when testing for correlations between genetic values for phenology versus observed collection dates). In both (B) and (C) there is co-gradient variation, but in (B) the genetic values for phenology traits (colors) predict local variation in natural phenology (residuals from local mean collection date) while in (C) the genetic values (colors) do not predict local phenology (residuals from local mean collection date).

## Materials and Methods

### Natural genotypes

We first conducted an experiment on a set of 101 naturally inbred genotypes (“ecotypes”) selected based on known collection dates either from the herbarium specimen label or recorded in the Arabidopsis Biological Resource Center https://abrc.osu.edu/ database. 47 of these ecotypes had not been studied in previous flowering time experiments and 75 had not been included in previous germination experiments. Seed was ordered from ABRC (94 accessions) or was germinated from herbarium sheets (7 accessions from Real Jardín Botánico Madrid, Komarov Botanical Institute, and Oslo Natural History Museum).

Seeds from herbarium specimens were cold-stratified in tap water at pH 7 and placed at 4°C for 7 d. Seeds were then directly sown into damp Fafard germination mix and grown in Conviron growth chambers under 14/10hr, 22/18°C days/nights. Four herbarium accessions germinated using this protocol. In a second round of germination, we surface sterilized seeds using standard protocols and then cold-stratified, as above. We plated these sterilized seeds onto MS+Gamborg’s vitamins + agar plates containing 1% sucrose and 10uM GA4 and then placed them in growth chambers, as above. Three additional herbarium accessions flowered using this protocol; these were transplanted to Fafard Germination mix. To induce flowering for seed collection, plants from both rounds of germination were exposed to 30d of 4°C with 10/18hr day/night cycles.

### Flowering and germination experiments

Ecotypes were grown in common conditions prior to the flowering time experiment, and flowering time replicates were descended from a single mother plant. For each ecotype, three replicates were grown in separate pots. Seeds were stratified at 4°C for 5 days before sowing in pots. Each pot was thinned to a single individual after the emergence of the second set of true leaves. Plants were grown at 22°C under 16h days of fluorescent light in a walk-in Conviron growth chamber (model MTPS144). Day of bolting and day petals appeared were both recorded as measures of flowering time.

Seeds from each replicate maternal plant in the flowering time experiment were collected and stored separately in dry conditions until the germination trial. Non-stratified seeds were kept in dry boxes at room temperature. For each treatment, forty seeds from each parent plant (or as many seeds as were available for replicates with low fecundity) evenly divided across 2 plates were sown on filter paper in petri dishes and germinated at 23/18°C during day/night with constant 16h daylength in a Conviron growth chamber. In total, 1,752 plates and >50k seeds were assayed.

Seeds were subjected to cold stratification at 4°C in the dark for 3 treatment lengths: 2 weeks, 3 days, and 0 days. Cold stratification can break primary dormancy, however 2 weeks of chilling can prompt secondary dormancy in the seedbank (Penfield and Springthorpe 2012). The difference in germination rate between 3 days and 0 days of stratification can therefore indicate primary dormancy while the difference in germination rate between 3 days and 2 weeks of stratification may indicate secondary dormancy. As a caveat, lower germination after 2 weeks of stratification could be due to other, unmeasured negative effects on germination, such as bacterial infection. We staggered the planting so that all the plates came out of stratification on the same day. The number of seeds that had germinated in each plate was recorded on days 1, 3, 5, 10, 14, 21, and 28. Seeds were considered germinated if the radicle was visible. Because maternal plants flowered at different times (and seeds were harvested when maternal plants senesced), germination rates for this experiment may be influenced by the length of the time between flowering and planting and any after-ripening that may have occurred. We also tested for these maternal effects among individuals of the same inbred line (Supplemental Material).

### Phenology from published experiments

We searched the literature for experiments on Arabidopsis that measured flowering time or germination traits across different natural inbred lines (commonly referred to as Arabidopsis “ecotypes”). The minimum number of ecotypes in any single experiment was 17. Our final set used data from 38 previous studies (31 included some measure of flowering time, 15 included germination, Table S1) that, combined with our new experiments described above, included over 3,000 ecotypes for 86 flowering time experimental conditions and 66 germination experimental conditions, although all ecotypes were used in only a subset of the trials and only 291 ecotypes had a reliable date of original collection from the wild.

We used this dataset of phenology measurements from the literature to create an estimate of genetic variation in flowering time and dormancy among ecotypes. We sought to gain statistical power by combining data from different experiments. To make phenotypes comparable, we standardized across treatments and experiments, transforming each flowering time experiment such that the earliest flowering accession had a value of 0 and the latest flowering accession had a value of 1.

We averaged standardized experimental flowering times across experiments, keeping vernalized, non-vernalized, and field experiments separate. Here, we use ‘vernalization’ to describe extended cold treatments applied to rosettes. Non-vernalized growing conditions uncover genetic variation in flowering time due to vernalization requirements that is masked under vernalized or fall-sown field conditions and may be important for determining phenology and life history in the wild (Wilczek et al. 2009). However, Arabidopsis plants growing in many natural settings are expected to experience changes in temperature and photoperiod that would be more similar to vernalized and field experiments (Li et al. 2010). By keeping the three treatments separate, we could test whether non-vernalized flowering time or vernalized flowering time was more predictive of wild phenology. Field flowering time was handled in two distinct manners: first, aggregated across all seasons, which may lead to less coherent estimates of field genetic flowering time if seasonal differences lead to meaningful differences in flowering time, and second, broken down further into spring (4 treatments over 2 experiments), summer (4 treatments, 1 experiment), and fall plantings (23 treatments, 9 experiments). To control for regional bias, this aggregation was performed both for all experiments and without experiments that tested only ecotypes collected from a single country.

Data collection approaches for estimating germination rate and dormancy were more heterogeneous than for flowering time. Depending on the experiment, dormancy was reported as the number of days after planting until a set percentage of germination was reached, the percentage of seed germinated a given number of days after planting, the number of days of storage until a set percentage of germination, or the germination rate after a given number of days of storage. Because of the difference in metrics across experiments, dormancy values were standardized by rank within each experiment with zero indicating low to no dormancy and one indicating high dormancy. Measures that reported a percentage germinated were ranked in the opposite direction from measures that reported number of days until germination or the number of days of storage before 50% germination. We found that standardized rank-based metrics that increase with dormancy (such as days to 50% germination) were correlated with rank based on metrics that decrease with dormancy (such as percentage of seeds germinated after a set number of days, Pearson’s *r* = 0.485).

These experiments captured variation in primary dormancy, or recalcitrance to germinate immediately after harvest. Secondary dormancy, or dormancy induced when a seed experiences conditions unfavorable to germination, has been studied in experiments that measured an increase in dormancy during storage. Primary and secondary dormancy could lead to different phenological and ecological outcomes (Martinez-Berdeja et al. 2020), so we averaged across experiments that measured primary or secondary dormancy separately to estimate each of these two traits. Secondary dormancy was not included in the Generalized Additive Models described below. Maternal conditions may be an important source of phenological plasticity; thus we also used our dormancy experiment to examine how variation in flowering time among maternal plants influences germination (Supplemental Material).

### Wild phenology

The date of collection (for ecotypes maintained by ABRC) or collection date (for ecotypes grown from a known herbarium record) was used as an estimate of reproductive phenology of individuals in the wild (Primack et al. 2004; Davis et al. 2015). We excluded records from regions where Arabidopsis has been recently introduced, like North America and Japan. Accessions from outside the native range or collected after the 320^th^ day of the year (which we deemed likely errors based on location, sometimes due to intentional plantings, and were greater than 2 standard deviations from the mean) were removed. Our geographic limits also excluded island accessions to the south of the Mediterranean, e.g. Cape Verde Island.

We hypothesized that genetic differences in wild phenology may be more apparent if we account for spatiotemporal environmental fluctuations causing plasticity. Therefore, we calculated photothermal units (PTUs) for each accession from the day of collection using monthly climate time series data from CRU (Harris et al. 2014), beginning with January 1 of each year, following the methods of (DeLeo et al. 2020) and (Burghardt et al. 2015). PTUs integrate the temperature and light experienced by a plant at a given location through the growing season and therefore may capture environmental cues relevant to phenology and better describe genetic variation in development (Wilczek et al. 2009; Brachi et al. 2010). We only sought to use PTUs to estimate how far into growing seasons plants were collected; we did not attempt to estimate PTUs since germination since germination timing was unknown. The models described below use the square root of PTU because the resulting distribution was closer to normal than the log transformation.

### Statistical comparison of collection date in the wild versus phenology in experiments

*H1: Range-wide comparison of phenology genetic values vs. collection date* We tested if genetic variation in normalized phenology measured on naturally inbred lines explained variation in the wild phenology of the parent of the line (Figure 2), using linear regression between the normalized phenology genetic values (flowering time and dormancy rank) and collection day. Vernalized, non-vernalized, and field flowering times were modelled separately, because the genetic variation in phenology uncovered by each treatment could relate to wild phenology in different ways. Under the hypothesis that vernalization and field conditions better recreate environments that a plant would experience at their geographic origin, these genetic flowering time measures should be more positively related to wild flowering time. Likewise, non-vernalized flowering times may better recreate the original temporal niches of summer annuals and thus be positively related to flowering time in these ecotypes. However, it is also possible that long non-vernalized flowering times indicate obligate winter annuals. In these plants, non-vernalized flowering times would be negatively related to wild phenology since later flowering times in non-vernalized experiments would indicate plants that overwinter and are collected early the next year (Figure 2).

We also investigated how range-wide differences in wild phenology related to regional differences in genetic trait values using a larger sample size, taken from a broader collection of ecotypes than just those with documented collection dates. This comparison included stock center ecotypes with experimental phenology data and known location-of-origin, but no recorded date of wild collection, and herbarium specimens for which we had not germinated seed and grown plants to measure traits in experiments. To do so, we first estimated wild flowering time for each stock center ecotype in experiments using herbarium records near the ecotype collection location. These wild flowering times were estimated from a previously published generalized additive model (GAM) with spatially-varying intercepts of herbaria collection dates from 2,655 Eurasian Arabidopsis records used in (DeLeo et al. 2020), which included year of collection as a nuisance variable, as collection date has changed over time across the range of Arabidopsis (DeLeo et al. 2020). Values of the smooth intercept surface were extracted at the coordinates of stock center ecotypes having an experimentally measured flowering time. These estimated wild flowering times were regressed against genetic values for flowering times under vernalized, non-vernalized, and field experiments and rank primary dormancy. In addition, for these same data we performed Spearman rank correlation between estimated wild flowering times and phenology genetic values.

### H2: Range-wide comparison of phenology genetic values vs. PTU

To account for plasticity in phenology due to differences among locations in the timing and progression of growing seasons, we also tested the relationship between phenology genetic values and the estimated PTUs at the time of collection from the wild. We conducted the same analyses across the range of Arabidopsis as described in H1, except that the PTU at collection was used instead of collection date. Specifically, we conducted linear regression between normalized genetic values with PTU at collection and we also conducted linear regression between genetic values and the PTUs at collection of nearby herbarium specimens.

### H3: Local within region variation

Next, we tested for the effects of genetic variation in flowering time and dormancy on phenological variation *within* local regions in the wild. By using a model term to first account for geographic, *among* region, variation in mean flowering time, we aimed to isolate the remaining local, *within*-region variation (Figure 3). This analysis, in essence, asks whether genetic variation in individual phenological stages explains natural phenological variation within local regions. Generalized Additive Models (GAMs) allow for model parameters to vary smoothly across space and thus can capture spatially varying patterns, e.g. in local average collection date (Yee and Mackenzie 2002; Yee and Mitchell 2006). We built a GAM for the dependent variable of collection date (*Y*) at location *j* of individual *i* which included covariates of experimental flowering time (β_1_), dormancy (β_2_), elevation (β_3_) and a spatially varying intercept (*µ_j_*):

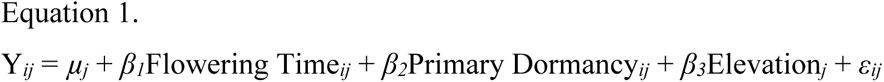

The smooth spatially varying intercept term *µ_j_* estimates spatial variation in mean collection date (e.g., due to environmental gradients). The upper limit on degrees of freedom for the spatially varying intercept was increased to 45 following the recommendations of Wood (2006). In models that allowed for higher degrees of freedom, the effective degrees of freedom did not meaningfully increase. Elevation was included because of its known importance to flowering times and spring onset (Vidigal et al. 2016; Gamba et al. 2023) and the high resolution of elevation data compared to smooth variation in GAM parameter surfaces. We fit GAMs using the ‘gam’ function in the ‘mgcv’ package in R (Wood 2006) using restricted maximum likelihood (REML), although Generalized Cross Validation returned similar estimates. Model fitting allowed for penalization of smooth terms to 0 so that uninformative covariates could be removed from the equation. Residuals were plotted using the ‘gam.check’ function in ‘mgcv’ (Wood 2006), and one ecotype (Nok-10) was removed from our analyses that was an extreme outlier based on its residuals. Spatial variation in *β* coefficients for flowering time and dormancy covariates (Eq 1) were not significant, so a model using a constant coefficient was used in our analyses. Field, vernalized, and non-vernalized flowering times were tested in separate versions of the model and tested different hypotheses with regards to wild phenology. To compare among the three measures of flowering time, a version of Equation 1 was fit on a subset of ecotypes that had all three flowering time measures, and Akaike’s Information Criterion (Akaike 1974) was compared.

In the wild, both flowering time and dormancy contribute to phenology. Thus, we also tested whether including interactions between flowering time and primary dormancy (*β_4_*) improved the model:

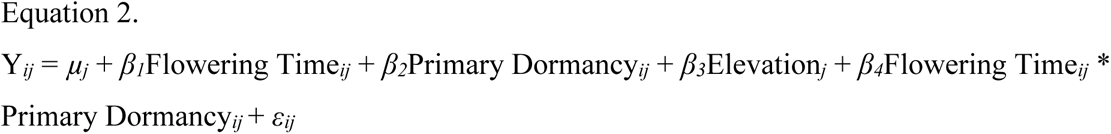

Finally, given the importance of plasticity in response to temperature in the timing of germination and flowering time, PTU might better capture genetic variation in phenology in the wild (Figure 2). Therefore, we also tested the models above using PTU in place of collection day to correct for climate.

## Results

### Genetic correlation among phenology traits in controlled experiments

Across all compiled experiments (both previously published and our new experiments) mean flowering times of ecotypes measured in vernalized and non-vernalized experiments were strongly positively correlated (Pearson’s *r* = 0.90, p < 0.001). Although fall-sown field trials might be expected to expose plants to cold seasonal temperatures that give vernalization cues, standardized flowering time genetic values in both non-vernalized and vernalized experiments were similarly correlated to published field experiment values (*r*_non-vernalized_ = 0.77, *r*_vernalized_ = 0.72, p < 0.001 for both). Primary and secondary dormancy were negatively correlated, but not significantly so (*r* = -0.15, p = 0.11). Despite known interactions between flowering time and dormancy due to seed maturation environment and pleiotropy of causal loci, only non-vernalized flowering time was significantly correlated with primary dormancy (*r* = -0.17, p = 0.02, Figure 4; for a heatplot of all phenotypes, see Figure S1). These correlations were qualitatively unchanged when using an alternate standardization method across experiments (Figure S2).

**Figure 4.**
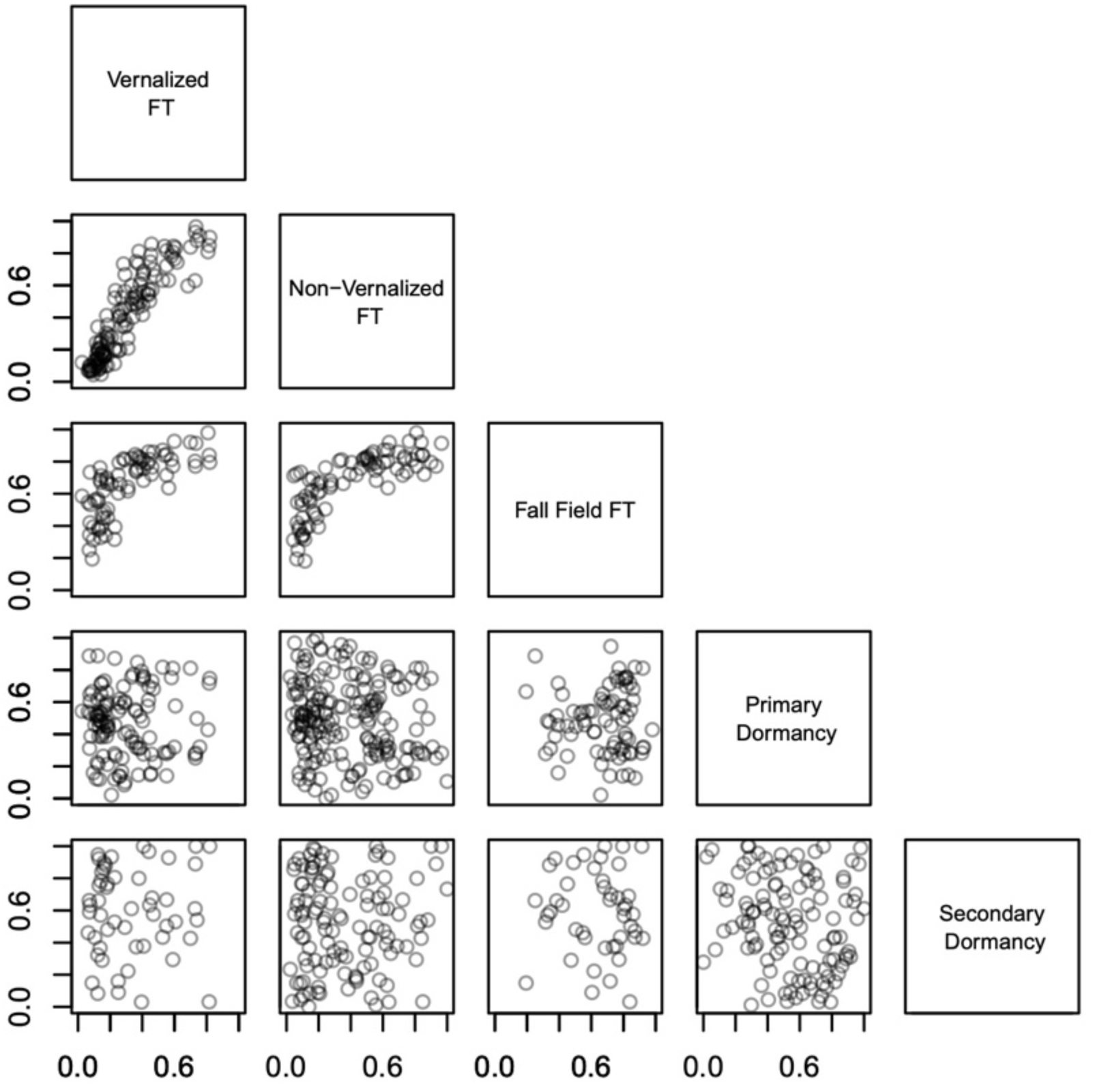
Standardized phenology traits combined across multiple published controlled experiments combined with our new experiments. Standardized flowering times were correlated across all three types of treatments (vernalized, non-vernalized, field), but they were weakly correlated with dormancy. See Methods for details on standardization method.

### H1: Comparing wild phenology with genetic values in experiments across the range

Experimental genetic values for phenology traits were modest predictors of collection day in simple linear models, with the only statistically significant predictors being non-vernalized flowering time (estimated slope = 27.7 days/standardized flowering time, r^2^ = 0.09, p < 0.001) and primary dormancy (-17.1 days/standardized rank dormancy, r^2^ = 0.02, p = 0.02). Flowering time genetic values under vernalized or field conditions did not significantly predict day of collection with a linear model (p > 0.05) (Figure 5). Flowering time and germination measured within individual experiments did not predict wild collection day better than our averaged genetic values, suggesting some power was gained by combining individual experiments into standardized values (Figure S3). These correlations were qualitatively unchanged when using an alternate standardization across experiments and when excluding herbarium specimens (Figures S4–S6).

**Figure 5.**
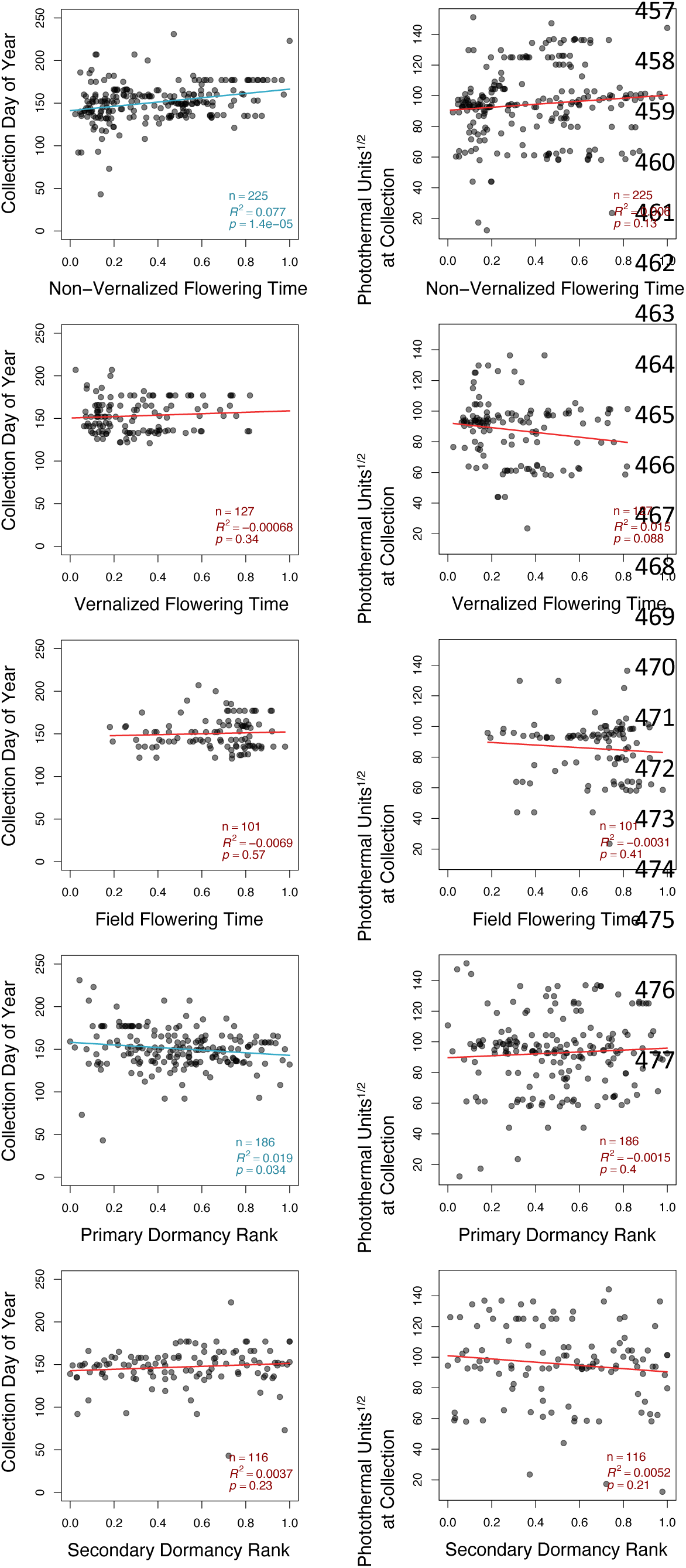
Date of collection (left panels) and PTUs (photothermal units) at collection (right panels) of wild plants (y-axes) compared to standardized genetic values for phenological traits from experiments (x-axes). Collection day of wild plants was positively related to non-vernalized flowering time in experiments. Collection day was negatively related to primary dormancy. The values shown for field flowering time experiments were calculated from fall-sown experiments only.

### Flowering time and dormancy genetic values predict regional variation in collection date of nearby herbarium specimens

Our model of collection date, above, was limited to the 227 ecotypes with both a known collection date and experimental phenology data. To expand our predictions across more of the Arabidopsis range, we tested whether genetic variation in flowering time in experiments predicts local mean collection dates of herbarium specimens across the landscape. First, we found that local mean collection dates across a landscape from >2500 herbarium collections were positively correlated with actual collection days of individual stock center ecotypes (which were not used in fitting the model, ρ = 0.51, p < 0.001 Figure S7), validating these local predictions of phenology in the wild based on herbarium collections.

As with the collection dates of the original maternal sources of stock center lines, genetic values for non-vernalized flowering times significantly but modestly predicted local mean herbarium collection dates (r^2^ = 0.10, p < 0.001). For comparison, latitude was a better predictor of mean herbarium collection date (r^2^ = 0.37, p < 0.001). These patterns were similar for vernalized (r^2^ = 0.07, p = 0.002, Figure 6), non-vernalized (r^2^ = 0.10, p < 0.001), and field measurements (all seasons grouped, r^2^ = 0.02, p = 0.13, although not significant in this last case). These results show that range-wide genetic variation in phenology measured in controlled environments likely corresponds to actual phenology of plants in nature. However, because plants in nature from different regions experience different environments, these patterns cannot determine the degree to which genetic variation causes the variation in natural phenology as opposed to genetic variation being correlated with environmental variation that causes phenological plasticity in nature (i.e. co-gradient variation, Figure 1).

**Figure 6.**
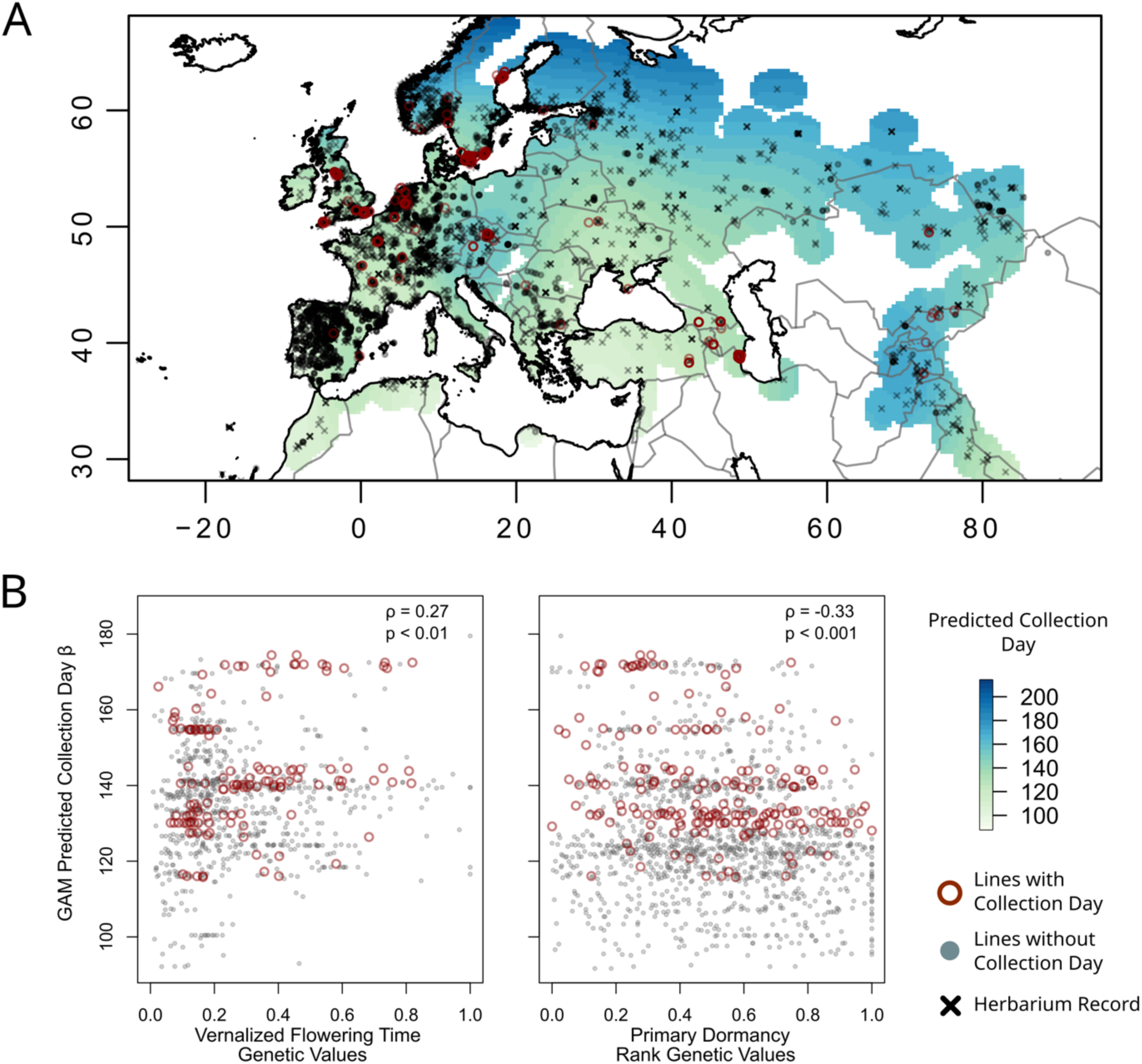
Genetic variation in flowering time experiments predicts collection dates of nearby herbarium specimens. A) There is geographic variation in collection dates as shown by estimated mean date from GAMs with spatially varying means, with earlier collections (light green) around the Mediterranean (DeLeo et al. 2020). B) Local mean collection dates were positively correlated with genetic variation in vernalized flowering time experiments (p < 0.01), and all ecotypes from locations with earlier mean collection dates in the wild (largely Mediterranean) had rapid flowering genetic values. Primary dormancy genetic values were negatively correlated with local mean collection dates (p < 0.001).

### H2: Range-wide comparison of phenology genetic values vs. Photothermal Units

Because environmental differences among locations can influence phenology (Fournier-Level et al. 2013), we also calculated PTU at collection and compared to these values to traits from experiments. However, PTU at collection was not significantly predicted by any phenology trait genetic values (Figure 5). Thus, while ecotypes with later flowering time genetic values tend to be collected later in the year, these later flowering ecotypes are not collected at higher PTU, i.e. later in the local growing season. The fact that phenology genetic values are correlated with date, but not PTU at collection, is consistent with the hypothesis that geographic climate variation maintains genetic clines in flowering time and germination, while plastic acceleration of flowering in warmer conditions (Figure 1) also contributes to the breeding value-collection date correlation (co-gradient variation).

### H3: Variation in phenology genetic values does not predict local, within-region variation in wild collection dates

We next examined whether genetic values could explain variation in natural phenology *within* regions (Figure 3). We used GAMs that included spatially varying intercepts and an elevation covariate to account for the among region variation in environmental effects on collection dates, leaving local variation to be potentially explained by genetic variation. However, we found that normalized flowering time (slope of non-vernalized flowering time = -8.2, p = 0.098) and dormancy (slope of dormancy = 2.2, p = 0.51) were not significantly related to day of collection in this GAM (Equation 1), although elevation effects were significant (slope of elevation = 0.012, p = 0.049, Figure 7, Table S2). In comparing models using vernalized, non-vernalized, and field experiments, we found that standardized non-vernalized flowering times led to lower AIC compared to other flowering time measures but higher AIC than a model without flowering time at all, although the difference was slight (difference < 2 for an AIC of 530). We used non-vernalized flowering times in the final models, in part because there were more ecotypes with non-vernalized flowering times (184 for non-vernalized vs 113 for vernalized or 78 for field).

**Figure 7.**
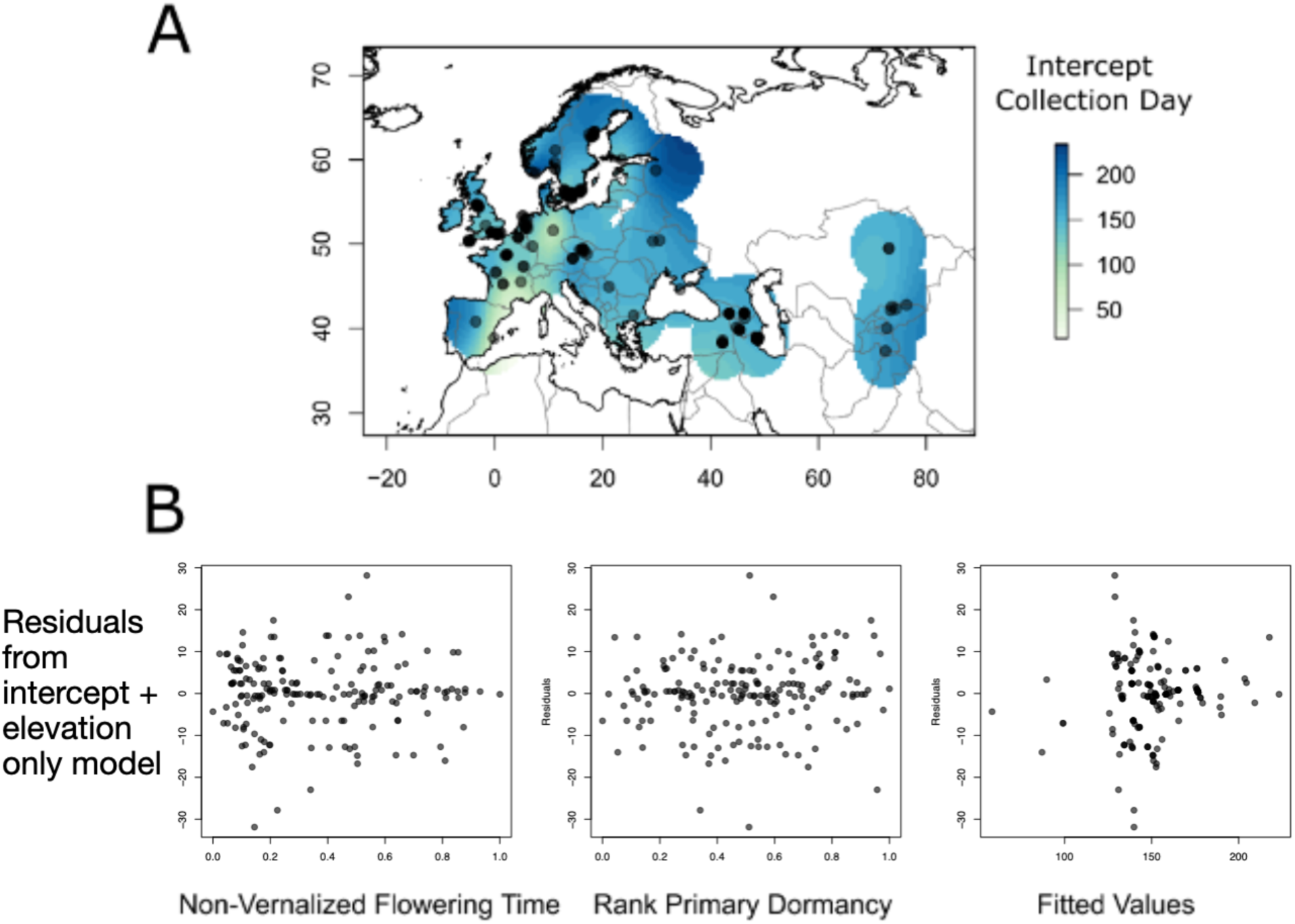
Top: Spatial variation in date of collection for 227 ecotypes with known collection dates. A) Color represents the intercept value for collection day at a location after fitting a linear (i.e. non-spatially varying) relationship with elevation, experimental primary dormancy and non-vernalized flowering time. Much of the variation in collection date was explained by the spatially varying intercept rather than being explained by experimental flowering time. B) Residuals of a GAM predicting collection date using only intercept and elevation parameters, compared to non-vernalized flowering time, primary dormancy, or fitted collection dates.

Although we found that flowering time, dormancy, and the interaction between the two were not significantly related to collection day, including phenology genetic values did lower the AIC and slightly increase the deviance explained by the model. This suggests that dormancy and flowering time could be related to variation in phenology within regions. However, most of the variation was described by the spatially varying intercept –a model with only a spatially varying intercept had a deviance explained of 80.1% – which could account for geographic genetic clines in phenology and plastic responses to geographic climate gradients.

As with the linear models, we also tested whether genetic variation in phenology in experiments could explain local variation in PTU of collection date in nature. Again, we found that flowering time and dormancy were not significantly related to wild PTU at collection in GAMs with spatially varying intercepts. Elevation was negatively related to PTU, indicating that higher elevation plants were collected earlier in growing seasons, reflecting a potential winter annual strategy (Figure 1, slope: -0.018 √PTU/m, p < 0.01). Additionally, elevation was positively correlated with date of collection (slope: 0.012 days/m, p < 0.05).

## Discussion

Plant phenology comprises multiple traits that contribute to fitness. In Arabidopsis, flowering time and germination can vary independently to create a landscape of possible life histories across environments (Debieu et al. 2013; Marcer et al. 2018; Martínez-Berdeja et al. 2020). This variation in phenology is likely partly maintained by selection, given that the traits vary geographically in association with climate (Fournier-Level et al. 2011; Vidigal et al. 2016; Exposito-Alonso 2020) and biotic pressures (Lyons et al. 2015; Davila Olivas et al. 2017), that QTL show evidence of local adaptation (Gamba et al. 2023; Lasky et al. 2024), and that the timing of individual phenological transitions is often correlated with fitness in experiments (Korves et al. 2007; Stock et al. 2015). Standing genetic variation is necessary for the timing of developmental transitions to evolve in response to selection by local environments, but little is known about the heritability of phenology in natural, wild individuals. Furthermore, because multiple developmental transitions influence phenology, the effects of variation in the timing of one stage might be compensated by (or, in under-parameterized experiments, confounded by) variation in the timing of other stages. Given widespread evidence that genetic variation in life history is adaptive, we investigated to what degree experimentally measured genetic variation in Arabidopsis phenology predicts phenology of plants in the wild, based on collection dates of natural history records.

The influence of genotype and environment on flowering time and dormancy have been well studied experimentally in Arabidopsis. Yet, common garden and controlled environment experiments must be designed thoughtfully to highlight genetic differences between ecotypes that are relevant to selection in natural environments (Karrenberg and Widmer 2008). While experimental design has increasingly recognized the importance of field conditions to acquire a measure of phenology that is more representative of nature (Wilczek et al. 2009; Anderson et al. 2012; Brachi et al. 2013; Poorter et al. 2016), there has been little comparison of controlled experiments, field or otherwise, to phenology in wild, naturally cycling individual plants. We found that flowering time and dormancy genetic values are related to date of collection and capture variation in phenology primarily *among* populations in the wild, but explain little variation *within* regions/populations.

### Genetic variation in flowering time and dormancy predicts variation in wild phenology among regions across the species range

Across the species range, both primary dormancy and non-vernalized flowering time significantly predicted collection dates, though much variation remained unexplained. These phenological transitions are not independent of each other. As seen in individual experiments (Martínez-Berdeja et al. 2020), we found that the average genetic values for primary dormancy and non-vernalized flowering time were negatively correlated (Figure 4). Furthermore, dormancy is strongly affected by environmental conditions during seed maturation (Penfield and Springthorpe 2012; Huang et al. 2015; Burghardt et al. 2016). Thus, interaction between flowering time and dormancy in the wild could stem from both genetic correlations and environmentally induced interactions. Despite our expectation for interactions between germination and flowering traits, models of collection day that included flowering time by dormancy interactions had a higher AIC and did not explain more deviance than a model with both traits separately. Including interactions between flowering time and dormancy in our models did not help to explain collection date in the wild.

We found that predicted wild collection date was positively related to flowering time genetic values when we extended our analysis to predict wild flowering times from a smoothed surface of collection dates fit to herbarium records. In our linear models, average flowering time breeding value was more closely related to this estimated day of collection than was the actual date of collection of ecotypes in experiments. However, a large portion of phenological variation within local regions remained unexplained even with phenological genetic values. We attempted to avoid records that were unusually young or old by only using specimens that had both flowers and fruits; moreover, both our herbarium and seed collections spanned nearly the entire year (Julian days 5–350 for herbarium records, 43–346 for seed collections). Still, herbarium records are known to skew slightly towards earlier in seasons (though removing herbarium specimens from our analyses did not change results) (Daru et al. 2018), while seed collections must be collected long enough after the initiation of flowering to allow for the development of some mature seeds. A single observation during reproduction, common for natural history collection vouchers, cannot resolve the uncertainty around when plants begin their vegetative growth in the wild, making it difficult to describe phenology of the full life cycle. With better information and in climates where growing conditions are clearly defined, it may be possible to estimate germination dates and full life cycle phenology from herbarium specimens (Bontrager et al. 2025a). Nevertheless, we found the expected strong latitudinal cline in collection dates of Arabidopsis, going from March to late June across the study region, and we previously found that collection date accelerates by ∼one week for every 1°C spring warming (DeLeo et al. 2020), suggesting collection dates do reflect natural phenology.

Spring onset differs among locations, and so a calendar date at a higher latitude or elevation may be functionally earlier in the season than a lower latitude or elevation. PTUs represent how much of a growing season has passed by a given date by recording temperature and daylight above a threshold. Thus, plants collected at lower PTUs may be early flowering (developmentally) despite a later collection date. Higher PTU may indicate plants growing as summer or fall annuals that germinated later in the year. In that case, ecotypes with higher PTUs would likely be fast cycling, earlier flowering plants (Figure 2). While we did find a negative correlation between flowering time genetic values in field and vernalized experiments versus PTU, we see the opposite in non-vernalized experiments. This last observation is somewhat surprising, since we expected plants that flower later in the absence of vernalization to be more likely to grow as winter annuals in the wild and flower early in the spring (Figure 1). Higher PTUs in these presumptive winter annuals may suggest that plants in these locations regulate their life histories to flower later in the year than expected by temperature and daylight alone.

There is some evidence that later flowering ecotypes could in fact be more flexible in their flowering time relative to germination than early flowering ecotypes (Miryeganeh et al. 2018), which could hide the signal of early spring flowering we expected in late flowering Arabidopsis.

### Within regions, genetic phenology values do not predict phenology in the wild

Because Arabidopsis exhibits substantial local within-population variation in phenology (Jones 1971; Brachi et al. 2013; Alonso-Blanco et al. 2016; DeLeo et al. 2020; Arana and Picó 2025), we asked how genetic variation in phenology was related to phenological variation within local regions. This within-region phenological variation can have fitness consequences. Variation contributes to population persistence in variable environments, as when different germination behavior provides bet hedging in the seedbank (Cohen 1967; Gremer and Venable 2014).

We found that neither dormancy nor flowering time genetic values from experiments were significantly related to collection date when we accounted for geographic variation in local mean collections dates in a GAM. However, flowering time and dormancy did improve GAMs for collection date over a model that included only geographic location and elevation, suggesting that genetic variation in the timing of germination and flowering time does explain a minor fraction of the phenological variation within regions. Non-vernalized flowering time was negatively, non-significantly related to local variation in collection date, perhaps because in many locations, genotypes flowering later in experiments without vernalization are more likely to behave as winter annuals, flowering early in the spring before spring annuals.

In our models, there was very little difference between non-vernalized, vernalized, and field-measured flowering times in predicting collection dates, despite the expectation that field experiments recreate conditions similar to nature. However, data from field experimental trials were available for a smaller set of ecotypes than non-vernalized and vernalized indoor trials, and we lacked Iberian ecotypes with field experimental data and date of collection, limiting the usefulness of this measure across the range. Our model provided weak evidence that the genetic values for flowering time and dormancy explained within-region variation in wild collection day. However, our set of ecotypes with known collection day and experimental indoor flowering time numbered 258, potentially still too few to detect strong statistical significance given the noise arising from plasticity in responses to local environmental gradients.

## Conclusion

We demonstrated that experimental measures of genetic variation in the timing of individual developmental transitions in Arabidopsis are significantly, but modestly, correlated with phenological variation in wild plants. On one hand, the existence of a relationship suggests there is some relevance of phenotypes measured in experiments to those exhibited in nature. However, the weakness of the relationships (especially locally, within regions) suggests the existence of either a) complex interactions between developmental stages to determine phenology, (b) low heritability in nature, or (c) extensive rank changing genotype-environment interactions across microsites. Genetic differences in Arabidopsis phenology across the species range are often cited as evidence of the adaptive importance of phenology (Stinchcombe et al. 2004; Fournier-Level et al. 2011; Samis et al. 2012; Brachi et al. 2013; Exposito-Alonso 2020). Incorporating information on between-population and within-population phenology diversity helps to clarify how selection may be acting on phenology across the landscape and how influential plasticity is in the phenology of this annual herb. Yet, genetic flowering times among populations across the species range are associated with phenological variation in natural history collections, due to some combination of genotypic and environmental effects (and their possible co-gradient variation) (Jones et al. 2024). While phenological plasticity is often of large magnitude, it may not be enough to maintain fitness under environmental change (Zettlemoyer et al. 2024). Ultimately, controlled experiments on many plants have suggested phenology is under selection in nature, but understanding more subtle environmental differences and stochasticity may help to clarify the evolution of phenology and translate genetic values into reliable predictions with and between populations.

## Acknowledgements

We thank the herbaria that allowed sampling of seed from specimens for generation of natural inbred lines used in experiments: Real Jardín Botánico de Madrid, Oslo Natural History Museum, and Komarov Botanical Institute. We are grateful for assistance with experiments from P. Patel, C. Yim, J. Kizer, T. Xia, V. Meagher, K. Turner, and E. Bellis and for helpful discussions with L. Burghardt. We thank reviewers for their constructive comments. Funding was provided by NIH award R35 GM138300 to JRL.

## Author Contribution

JRL conceived of the project with TEJ, DLD assisted in experiments, VLD conducted experiments, and analyzed data, VLD and JRL led writing, all authors contributed to the interpretation and writing of the manuscript.

## Data Accessibility Statement

Original data will be included as supplemental tables. Prior studies from which phenology data was sourced will be listed in a supplemental table.

## Supplemental Material

### Plasticity due to maternal effects

Maternal conditions may be an important source of phenological plasticity; thus we used our dormancy experiment to examine how variation in flowering time among maternal plants influences germination. Variation in phenology may not be adaptive in regions where there is harsh seasonality or environmental conditions are less variable year to year (Cohen 1967). Similarly, observations of phenology in the wild could be temporally variable because of year-to-year environmental differences (Walker et al. 1995; Hu et al. 2017; Postma and Ågren 2018) or genetically diverse individuals germinating from the seedbank (Ratcliffe 1976). Among replicate maternal individuals in the same controlled experiment, variation in phenology is likely due to responses to very subtle environmental differences and some developmental stochasticity.

For each ecotype and for each maternal replicate, we used R packages ‘drc’ (Ritz et al. 2015) and ‘drcSeedGerm’ (Onofri et al. 2018) to fit a logistic function to germination counts to estimate three parameters: maximum germination proportion, time to 50% germination, and slope of the germination curve. A Generalized Linear Mixed Model was fit using the lmer package in R (Bates et al. 2015) to estimate the influence of relative flowering time of maternal replicates of each ecotype on germination traits. Because plasticity or responsiveness to environmental cues may confer greater fitness in some environments and not others (Alpert and Simms 2002; Baythavong 2011), germination traits and variation were regressed against maternal flowering time and location of origin. We performed hierarchical clustering of a distance matrix of phenotypic variation among ecotypes to group ecotypes that reacted similarly to different stratification treatments.

We found that some ecotypes had more consistent phenology across maternal lines. For example, DAM1 maternal replicates clustered together in germination across cold treatments, whereas Yeg-7 replicates did not (Figure S8A). Cold stratification altered expression of variance among maternal sources. In Yeg-7, for example, maternal lines P vs A had similar total germination following 3 days or 2 weeks of stratification but different germination for 0 days stratification. Thus, stratification (or lack of stratification) could contribute to diverse phenology for similar genetic backgrounds under natural conditions. The first and second principal components in an analysis of flowering and germination traits explained 24.55% and 17.85% of the variation respectively, but again maternal replicates of ecotypes did not always cluster in PC space (Figure S8B). The variance in phenology for an ecotype was not related to collection date or geographic origin (latitude and longitude, Figure S8C), suggesting that clinal variation in maternal plasticity effects do not confound our earlier large-scale analyses comparing trait genetic values with phenology in nature. Nevertheless, these effects may further obscure genetic contributions to phenology in nature.

We expected to see more variation among ecotypes from regions with favorable growing conditions year-round. However, the lack of a geographic association between variance in germination traits and flowering time among replicate individuals of an ecotype may indicate geographically consistent selection on the canalization of phenology.

**Figure S1.**
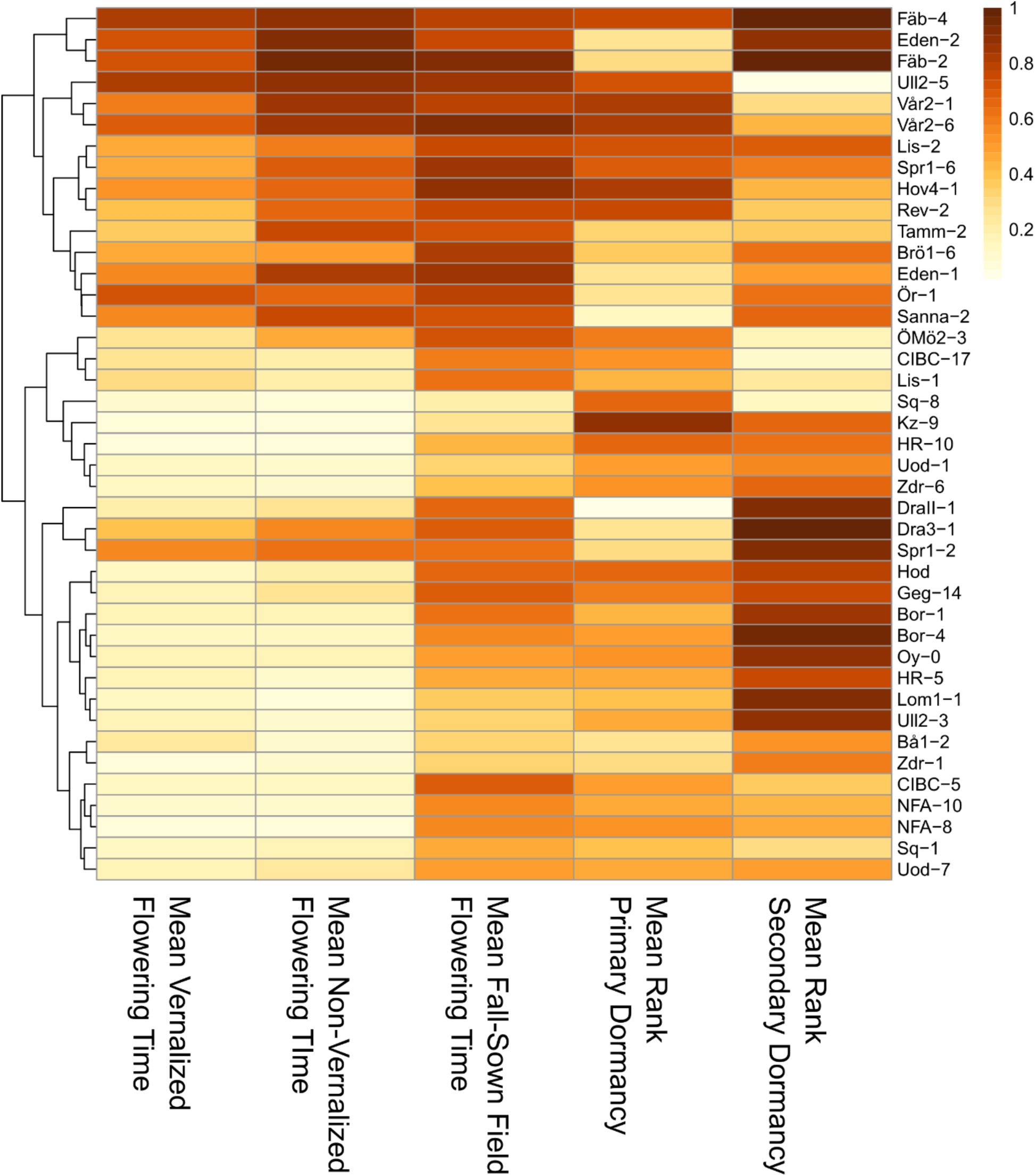
42 ecotypes have measures of vernalized, non-vernalized, and field flowering times as well as primary and secondary dormancy traits (standardized trait units are shown). The latest flowering accessions tend to be late flowering across all treatments, but primary and secondary dormancy vary independently from flowering time.

**Figure S2.**
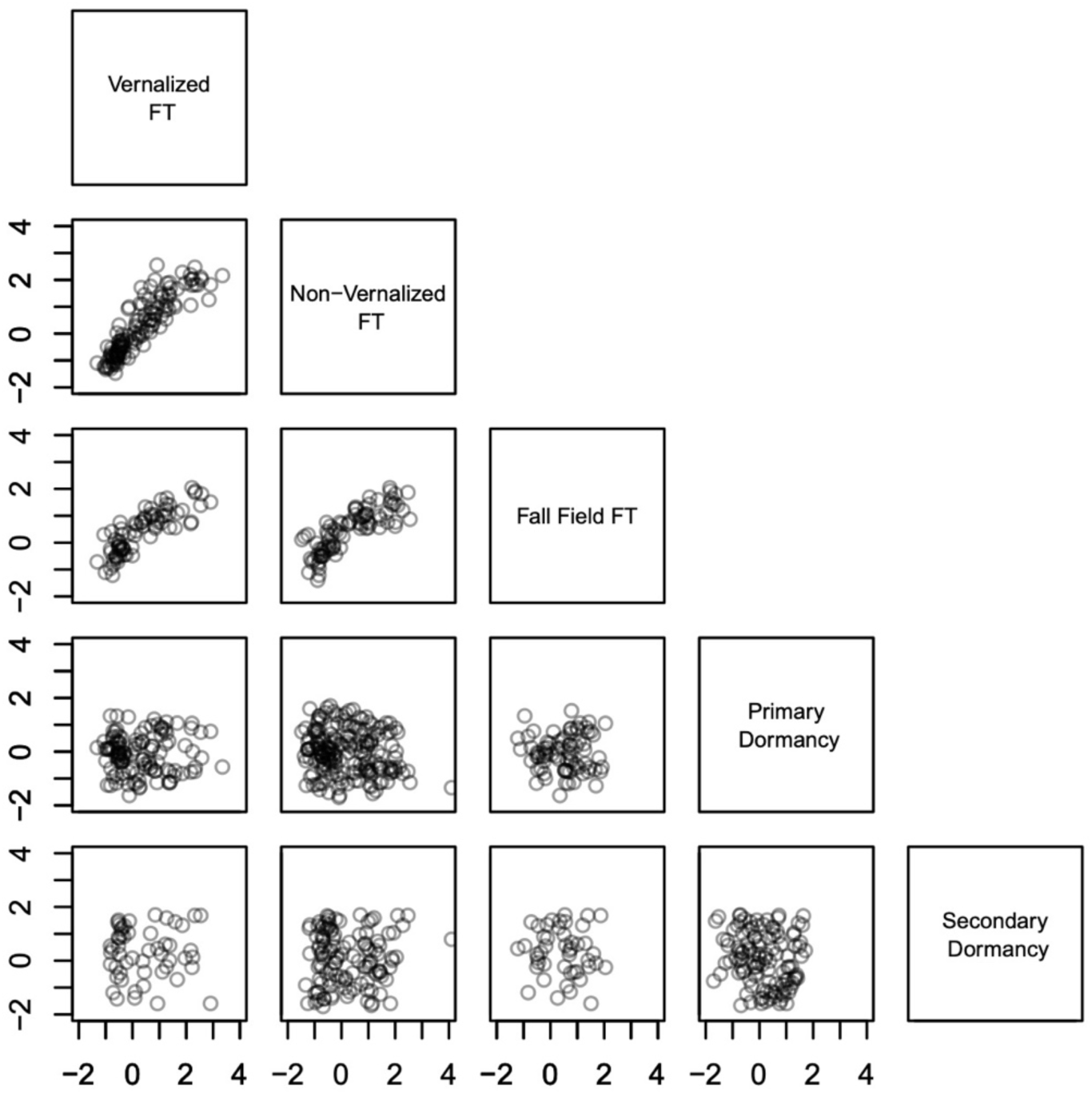
Standardized phenology traits combined across multiple published controlled experiments combined with our new experiments, using an alternate standardization (compare with Figure 3). Here, each experiment’s values were standardized to mean zero, unit standard deviation.

**Figure S3.**
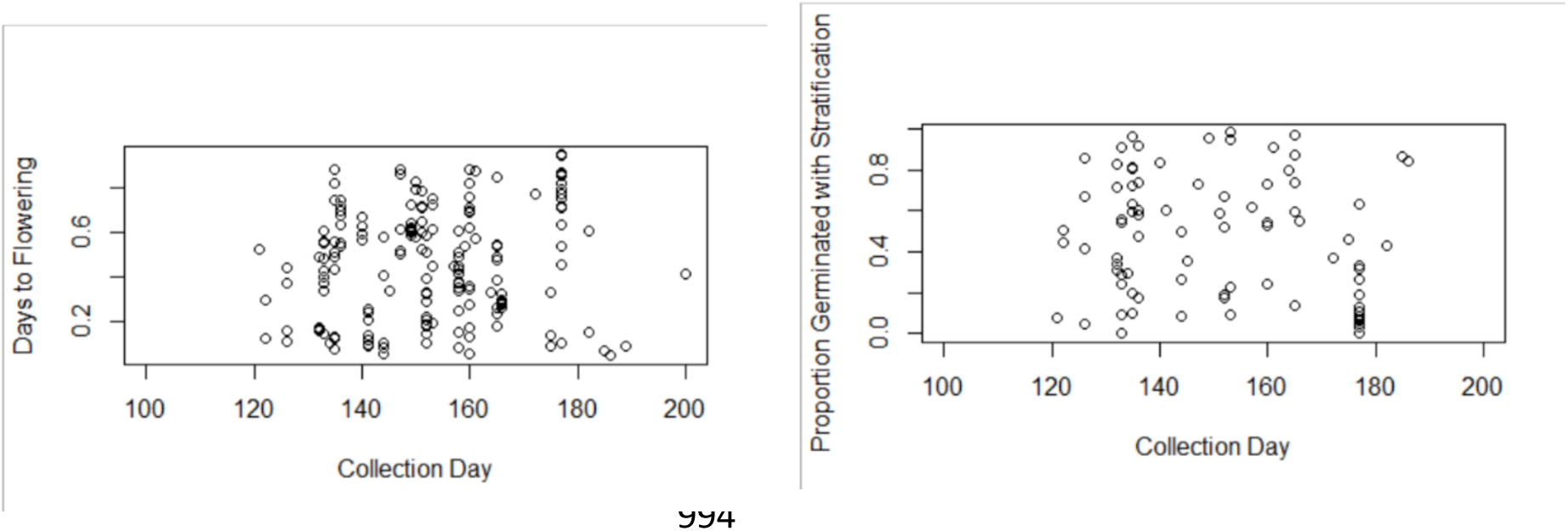
Flowering times and germination rates within experiments do not predict wild phenology better than average values across multiple experiments. Left, non-vernalized flowering times at 16°C (using the same criteria to remove collection day and geographical outliers as in the full set of accessions) Alonso-Blanco et al. (2016) did not significantly predict collection day (p > 0.2, n = 173). Right, germination rates after 4 days of stratification at 4°C Martínez-Berdeja et al. (2020) did not significantly predict collection day (p > 0.1, n = 84).

**Figure S4.**
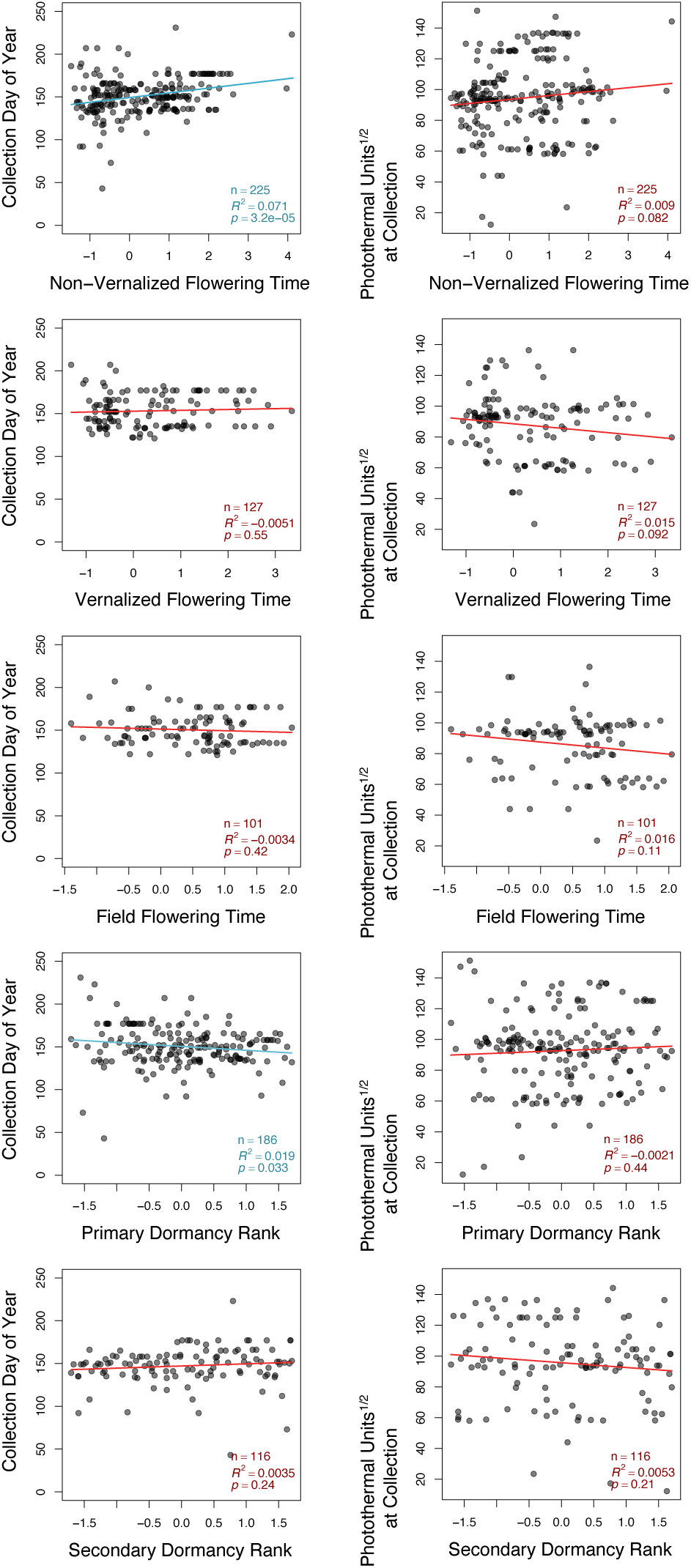
Date of collection (left panels) and PTUs (photothermal units) at collection (right panels) of wild plants compared to standardized genetic values for phenological traits from experiments, here using an alternate standardization (compare to Figure 4). Here, each experiment’s values were standardized to mean zero unit standard deviation. The values shown for field flowering time experiments were calculated from fall-sown experiments only.

**Figure S5.**
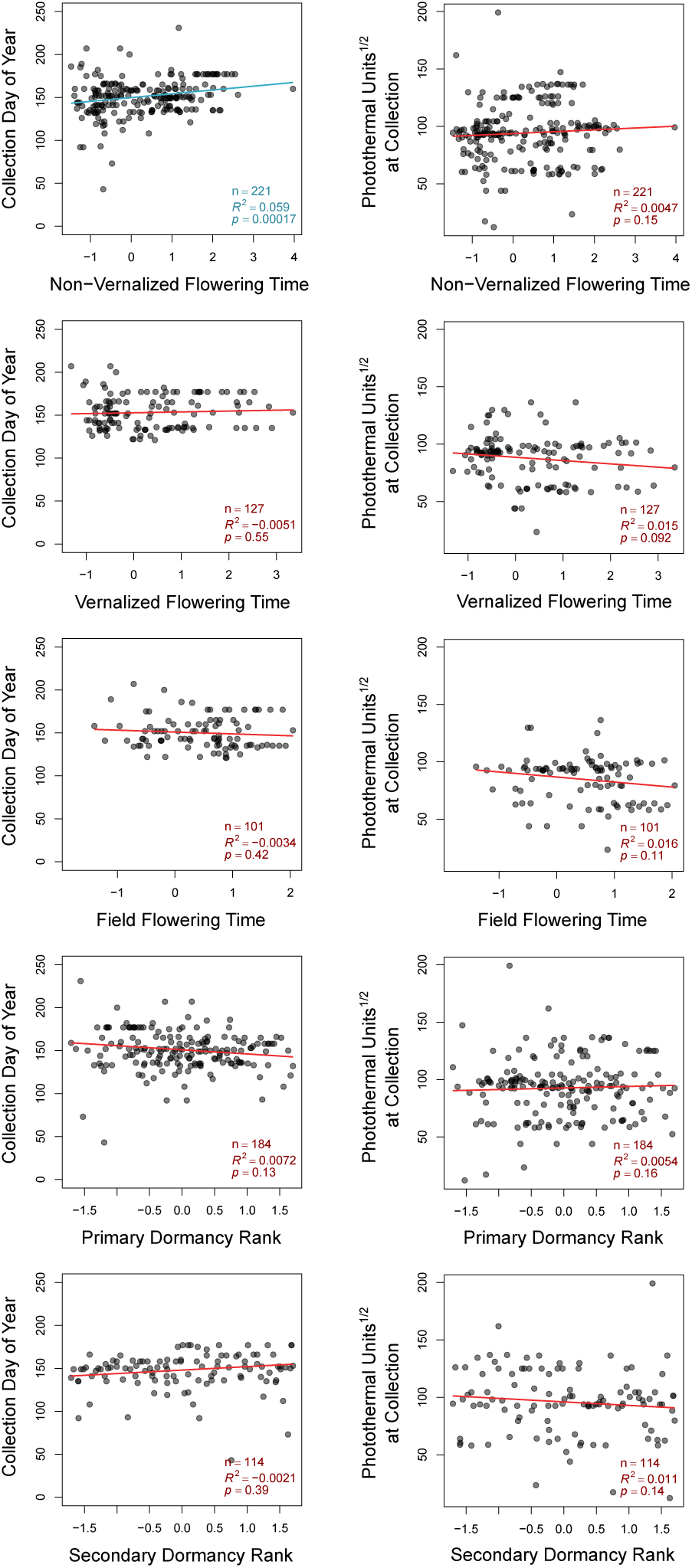
Date of collection (left panels) and PTUs (photothermal units) at collection (right panels) of wild plants compared to standardized genetic values for phenological traits from experiments, here using an alternate standardization and excluding herbarium specimens (compare to Figure 4). Here, each experiment’s values were standardized to mean zero unit standard deviation. The values shown for field flowering time experiments were calculated from fall-sown experiments only.

**Figure S6.**
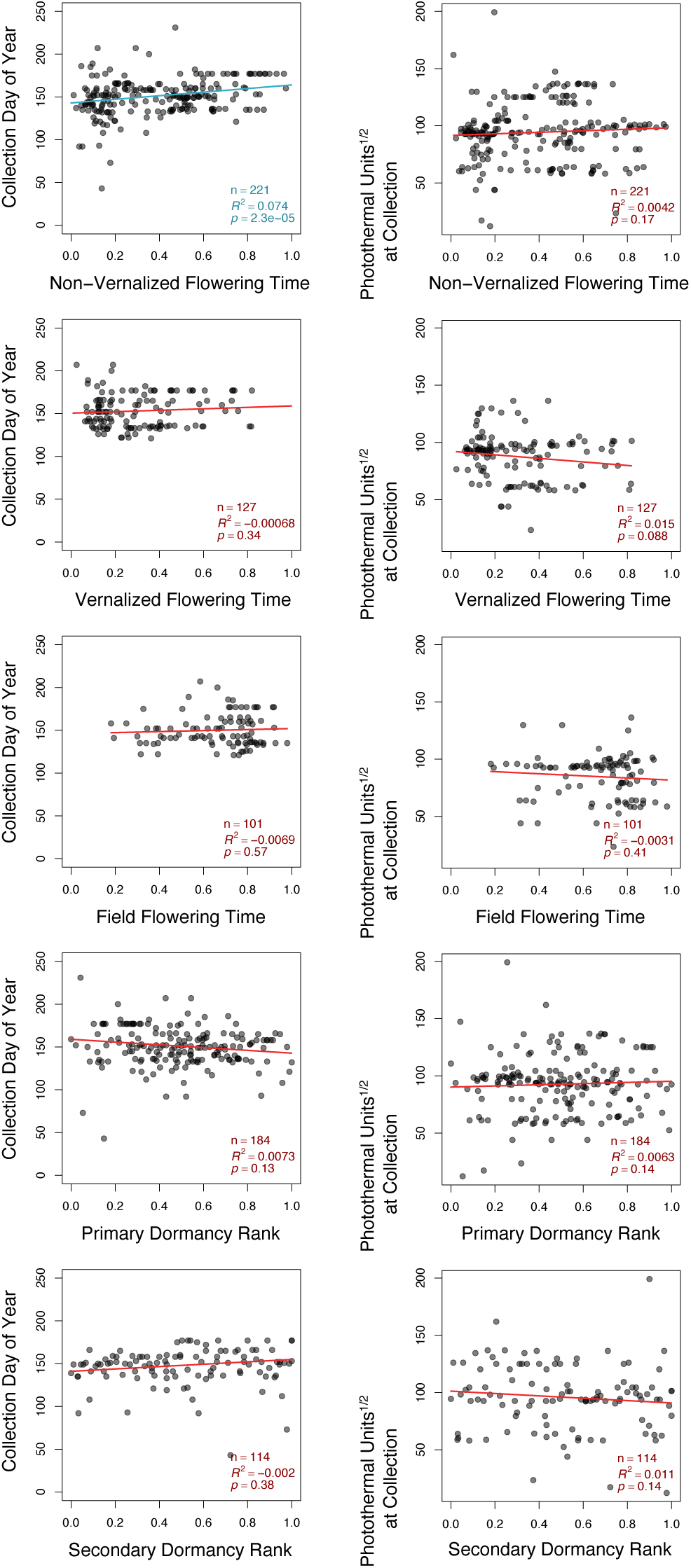
Date of collection (left panels) and PTUs (photothermal units) at collection (right panels) of wild plants compared to standardized genetic values for phenological traits from experiments, excluding herbarium specimens (compare to Figure 4). The values shown for field flowering time experiments were calculated from fall-sown experiments only.

**Figure S8:**
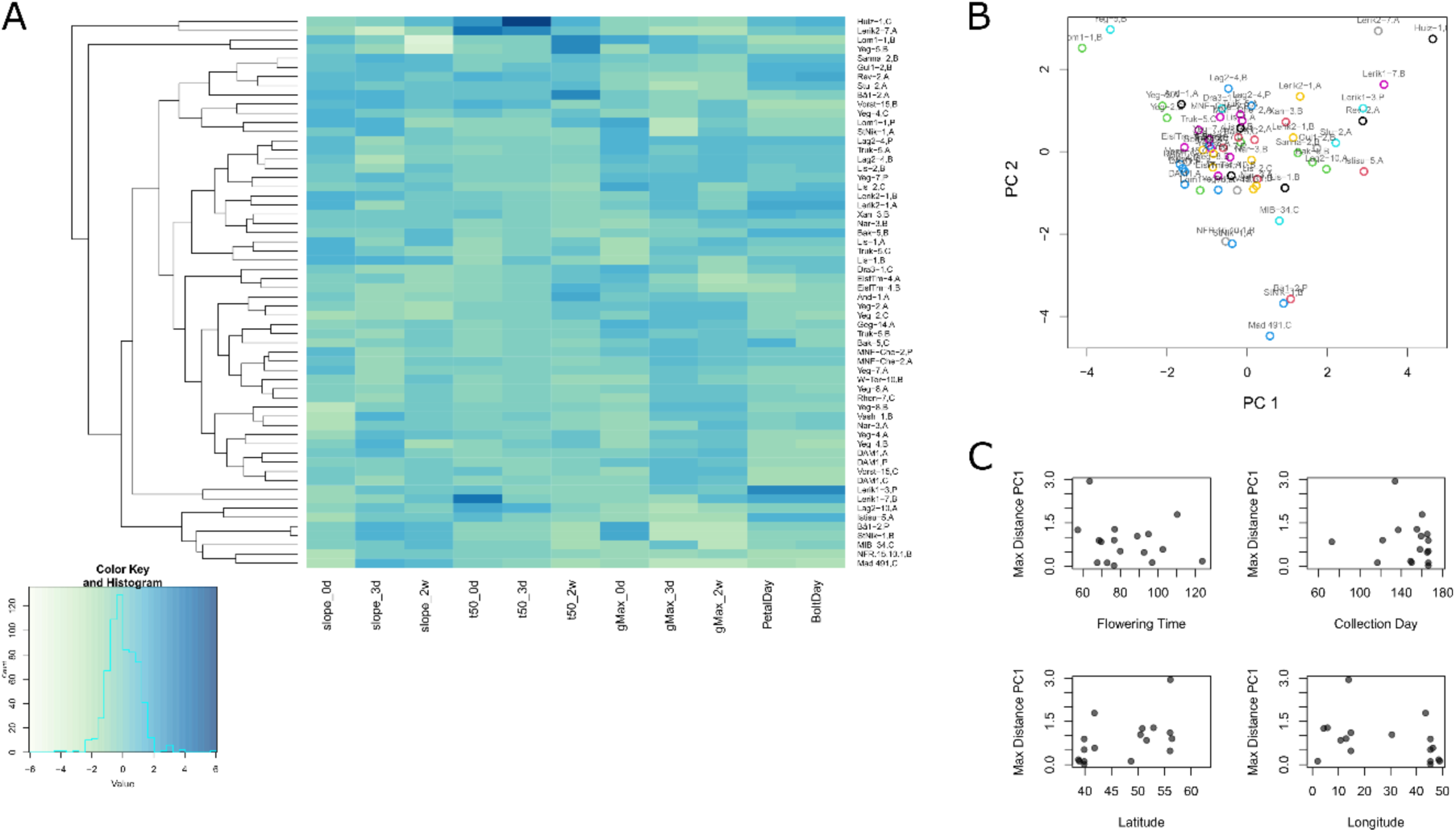
Variation in experimental phenology among maternal lines. A) Early flowering lines tended to have lower germination under 0 days stratification relative to 3 days or 2 weeks stratification, and later-flowering lines tended to have longer times to 50% germination. However, there was variation across traits. Replicate maternal lines of the same ecotype did not always cluster. B) Maternal lines of the same ecotype did not always cluster together in PCA of phenology traits. Lines belonging to the same ecotype are represented with the same color, although colors are not unique across ecotypes. PC1 was positively associated with time to 50% germination at 0 days and 3 days stratification and flowering time and negatively associated with time to 50% germination at 2 weeks stratification and max germination under all treatments. C. Flowering time, collection day, or geographic origin did not explain variance among maternal lines.

**Table S1.** [see attached supplemental Excel file]. Experiments included in our study and phenotypes represented.

**Table S2:** Model statistics for other variables in models of within-region phenological variation. P-values for the estimates are given in parentheses. Phenology genetic values did not significantly predict within-region variation in phenology.

| Model | Non-Vernalized Flowering Time | Primary Dormancy | FT*Dormancy | Elevation | Adj R<sup>2</sup> |
| --- | --- | --- | --- | --- | --- |
| Collection Date<br>~ $\mu$ + FT + Elevation | -3.25 (0.35) | NA | NA | **0.0148**<br>**(0.006)** | 0.883 |
| PTU<sup>1/2</sup> ~ $\mu$ + FT + Elevation | -1.21 (0.75) | NA | NA | **-0.0147**<br>**(0.006)** | 0.876 |
| Collection Date<br>~ $\mu$ + Dormancy + Elevation | NA | 1.98 (0.56) | NA | **0.0126**<br>**(0.04)** | 0.876 |
| PTU<sup>1/2</sup> ~ $\mu$ + Dormancy + Elevation | NA | 2.56 (0.48) | NA | **-0.0181**<br>**(0.009)** | 0.858 |
| Collection Date<br>~ $\mu$ + FT + Dormancy + Elevation | -8.19 (0.10) | 2.19 (0.51) | NA | **0.0120**<br>**(0.05)** | 0.881 |
| PTU<sup>1/2</sup> ~ $\mu$ + FT*Dormancy + Elevation | -5.41 (0.56) | 5.39 (0.43) | -6.86 (0.66) | **-0.0183**<br>**(0.009)** | 0.864 |

